# Mosquitoes asses interspecific competition by larvae-released semiochemicals during oviposition habitat selection

**DOI:** 10.1101/2024.03.31.587456

**Authors:** Nimrod Shteindel, Yoram Gerchman, Alon Silberbush

**Affiliations:** Faculty of Natural Sciences, University of Haifa, Israel; Oranim Academic College, Israel

## Abstract

Numerous species of animals alter their behavior in response to increasing competition. To do so, they must possess the ability to detect the presence and density of interspecific competitors. We studied the role of semiochemicals released by increasing densities of larval *Culiseta longiareolata* Macquart on female oviposition habitat selection in two sets of field mesocosms. Similarly to *C. longiareolata* larvae, subordinate *Culex laticinctus* Edwards are periphyton grazers who dwell in rain-filled pools in the Mediterranean region. We show that *C. laticinctus* females oviposited significantly less in mesocosm pools that were treated with crowding signals originating from *C. longiareolata* larvae. In a second field experiment, we placed a similar number of larvae directly inside the 50 L mesocosms. These low-density mesocosms did not affect *C. laticinctus* oviposition but were attractive to conspecific oviposition. These results increase our understanding of the female’s ability to detect species-specific signals indicating increased larval competition.

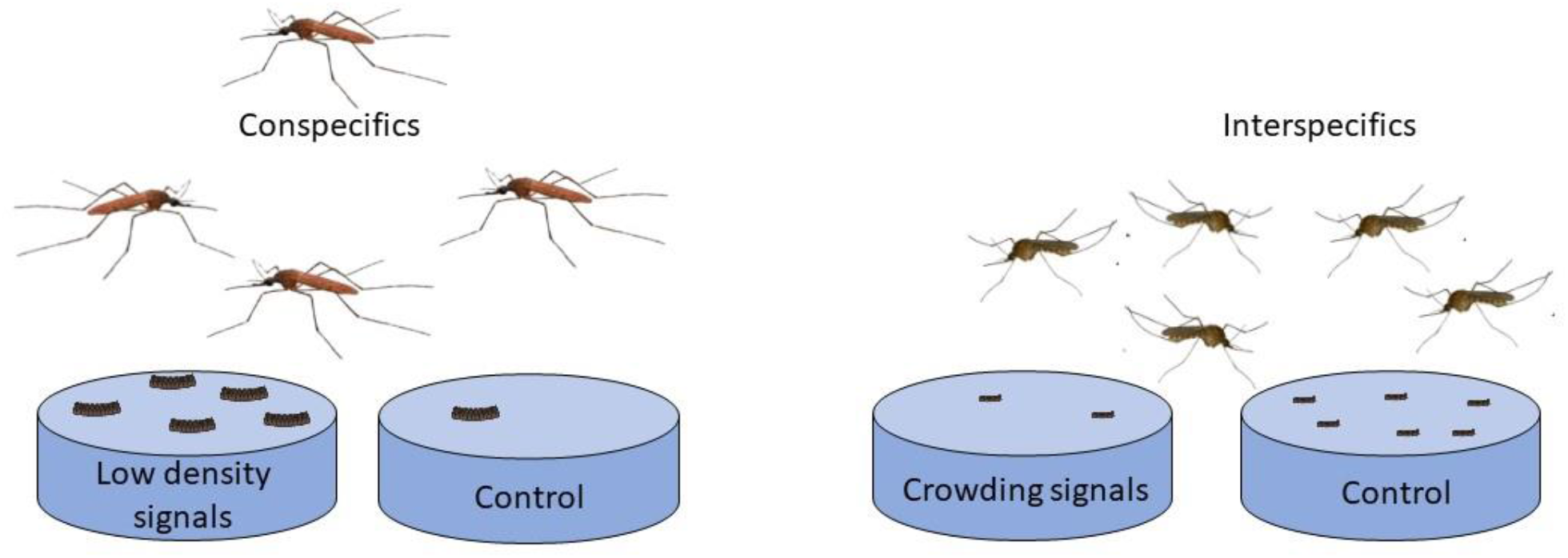

## Introduction

Interspecies competition, is a reciprocally negative interaction between populations of two or more species (Rockwood 2015, Begon et al. 2006). This fundamental interaction between species that share a similar niche is considered one of the most important factors in the shaping of the ecological community (Begon et al. 2006, Morin 2011), as well as an important driver of evolutionary speciation (Hutchinson 1959). The high cost of competition is associated with several alterations in animal behavior. In the presence of a competing species, individuals alter their foraging activity (Begon et al. 2006, Abrams 2010, Dhondt 2012), as well as produce aggressive interference and shifts in mating success (Dhondt 2012, Grether et al. 2017). Habitat selection is another mechanism affected by competition, and some animal species shift their activity, in space (to a less desirable habitat) or in time (to a different time of day or a different season) to reduce competitive interactions (Abramsky et al. 1990). These behavior alterations happen in response to increased densities of both populations and thus require the ability to identify competitor presence as well as an estimation of conspecifics (Rosenzweig 1981). Although several studies demonstrated the ability of competitors to do just that (Morris et al. 2000, Abramsky et al. 1990, Sandlin 2000), the mechanism of identification is usually overlooked.

Chemical signals or semiochemicals, are an important source of information for numerous animal species. This is especially true in aquatic systems since odorants tend to travel better in water compared to auditory and visual signals (Brönmark and Hansson 2012). Aquatic species alter their behavior in response to semiochemicals indicating a food source, predator presence, or conspecific density (von Elert 2012) . With few exceptions e.g. (Wu et al. 2019), studies on behavior alteration as a result of semiochemicals associated with interspecific competitors looked at mere recognition and overlooked the density-dependent effects.

Ovipositing mosquito females (Diptera: Culicidae) are excellent models for the study of semiochemical effects on habitat selection. Mosquitoes are characterized by a complex life cycle where adults are free to range the landscape but the immature are confined to the aquatic habitat where they hatched. Female mosquitoes provide little parental care beyond the selection of an appropriate oviposition site, making oviposition a critical factor in larval survival. Ovipositing females are attracted to several bacteria-released semiochemicals that are associated with nutrients for future larvae (Takken 1999, Bentley and Day 1989). Females also use semiochemicals to detect predators and avoid oviposition in sites where predation risk is high (Bentley and Day 1989, Angelon and Petranka 2002) . In addition, gravid females can quantify predators (Silberbush and Blaustein 2011), and conspecific larvae (Wasserberg et al. 2014), rather than simply be aware of their presence. In this study, we examined the ability of ovipositing mosquito females to detect and respond to the presence and density of semiochemicals originated by larvae of a dominant competitor species.

### Study species

*Culiseta longiareolata* Macquart is a highly abundant species throughout the Mediterranean region (Becker et al. 2010). The females typically oviposit in small, often temporary, rain-filled bodies of water and are often the earliest colonizers of these habitats following rain (Van Pletzen and Van der Linde 1981, Blaustein and Margalit 1996, Ward and Blaustein 1994). Because these rain-filled pools are both ephemeral and limited in number, they are a valuable resource to amphibians and aquatic insects. *Culex laticinctus* Edwards is a mosquito species whose larvae are often associated with *C. longiareolata* breeding sites (Margalit and Tahori 1974, Becker et al. 2010). The larvae of this species are considerably smaller in comparison to *C. longiareolata* larvae (Figure 1). *Culiseta longiareolata* larvae are considered herbivorous and feed mainly on periphyton algae and bacteria (Van Pletzen 1981). Nevertheless, fourth instar *C. longiareolata* larvae are considered as highly aggressive competitors of other freshwater species such as *Bufo virdis* tadpoles (Blaustein and Margalit 1994) and other mosquito larvae (Tsurim et al. 2013). This aggressive behavior towards other aquatic dwellers may result in the death of larvae of other mosquito species (Al-Saadi and Mohsen 1988, Shaalan 2012) as well as vertebrates such as *Bufo virdis* tadpoles (Blaustein and Margalit 1994).

**Figure 1.**
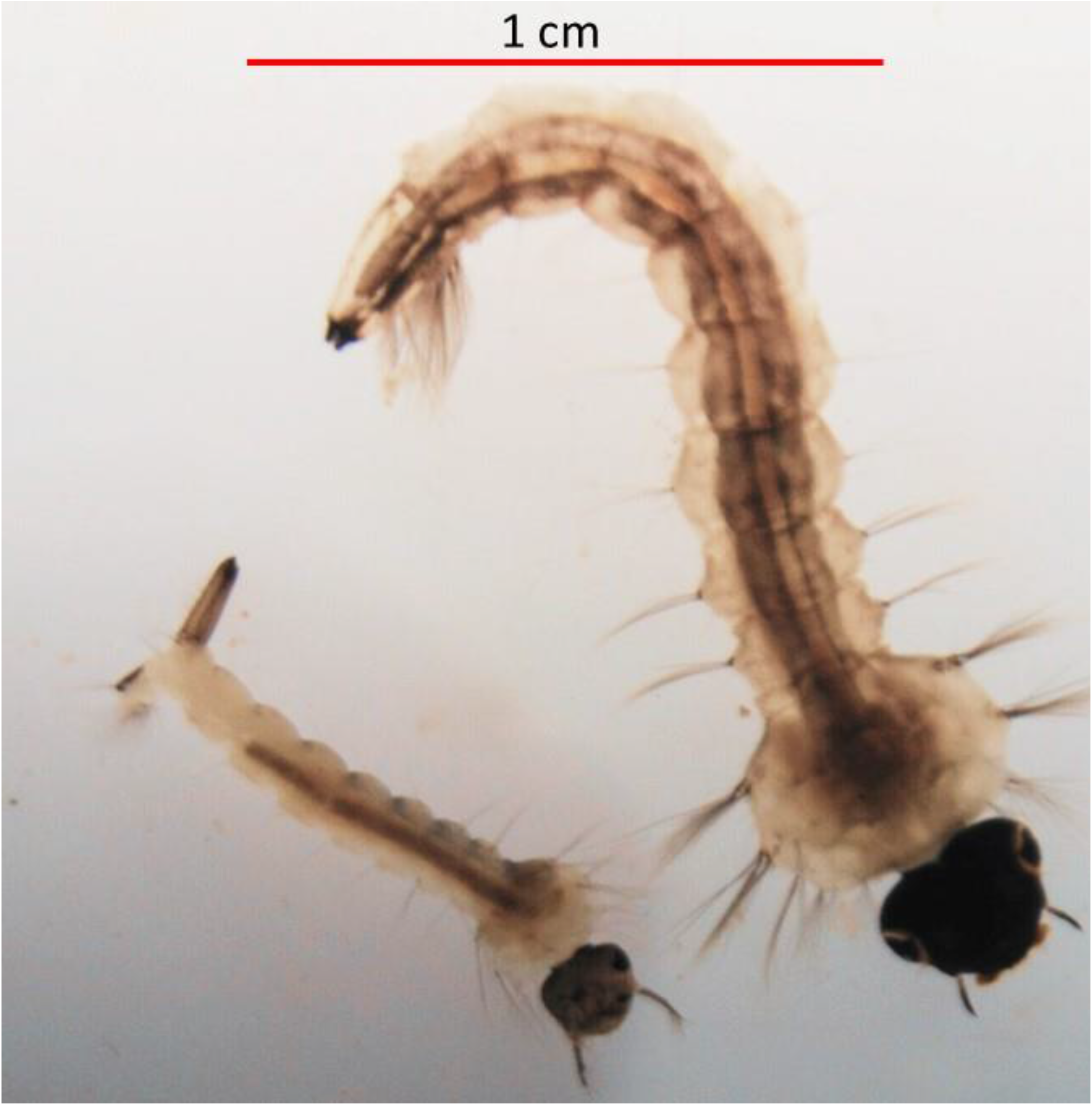
Fourth instar larval: *Culiseta longiareolata* (top) and *Culex laticinctus*.

The purpose of this study was to examine the effects of chemical signals produced by high densities of the dominant competitor *C. longiareolata* larvae on ovipositing mosquito females. The study was conducted with field mesocosms that mimic rain-filled pools that are the natural larval habitats of both species. We hypothesized that water with high *C. longiareolata* larval density will contain crowding signals. These signals, indicating high competition will be associated with larval habitat of poor quality. Ovipositing females are therefore hypothesized to avoid these habitats. In a second experiment, we examined the effects of the actual larvae who were not subjected to crowding. According to the Ideal Free Distribution theory, conspecifics should always prefer habitats with lower density (Fretwell and Lucas 1969). Subordinate competitor species are likewise hypothesized to avoid competition and reduce oviposition in habitats containing competitors.

## Methods

### Field experiments

Field experiments were conducted at the Oranim college campus botanical gardens (Tivon-Israel) between May and June 2023, a period that is peak activity for ovipositing mosquitoes in that area. We monitored mosquito oviposition in black plastic pools sized 66.04×50.8×15.24 cm. Pools were organized in a randomized block design, containing eight blocks of three treatments (n=24) randomly distributed within each block. Pools within a block were placed ∼1m apart and blocks were spaced ∼10 m from each other. Pools were filled with ∼50L tap water and supplemented with 10g of rodent chow to enhance oviposition. We collected mosquito egg rafts daily from the water surface of each pool. The collected egg rafts were hatched and the larvae were raised to the 4^th^ instar and identified to species using (Becker et al. 2010).

#### Crowding signals experiment

We produced water-containing signals of highly crowded *C. longiareolata* larvae by placing 200 4^th^ instar larvae in 400 ml plastic cups for 24 hours. This density of 500 larvae/L is considered high but not an untypical density for this species (Blaustein and Margalit 1996, Blaustein and Margalit 1994). A second treatment with medium crowding signals included 20 larvae in 400 ml (50 larvae/L), and a third set included control cups with no larvae. Larvae were fed with ∼0.05 grams of finely grounded fish flakes (42.2% crude protein) that were added to each cup. We filtered the water daily and replaced the dead and pupated larvae. Conditioned water (without the larvae) was added to the experimental pools each day at sunset. All pools were emptied and refilled every 5 days.

#### Low-density larvae experiment

We placed 2 densities of living *C. longiareolata* larvae directly in the field mesocosms. The higher density included 200 4^th^ instar larvae, roughly the amount larvae originated from a single egg raft (Van Pletzen and Van der Linde 1981, Al-Jaran and Katbeh-Bader 2001). The lower concentration treatment consisted of 20 larvae per pool, and the third pool was a control pool without larvae. Pupated larvae were replaced daily and all pools were emptied and refilled every 5 days.

### Statistical analysis

We used the total number of all egg rafts, per pool, collected for each mosquito species across all dates as a dependent variable. We used square root transformations of these values with an addition of 0.5 to all values, to homogenize among-treatment variance (Yamamura 1999). The homogeneity of variance was then tested using Levene’s test. We conducted separate univariate ANOVAs for each mosquito species in each of the experiments using “Block” and “Treatment” as fixed factors. Treatment means were compared using Fisher’s protected LSD when the main effect of treatment had p<0.1 using α = 0.05 for individual LSD comparisons. All analyses used SPSS statistics for Windows version 24 (IBM 2016).

## Results

The first field experiment ran for 15 days (May 3^rd^-May 18^th^ 2023) and the second for 25 days (May 22^nd^-June 16^th^ 2023). During these periods, we collected a total of 335 and 340 egg rafts from both setups respectively. One of the blocks contained an especially low number of egg rafts during the first period and was removed from the analysis. All egg rafts collected in both field experiments belonged to one of three mosquito species. *Culiseta longiareolata* (Macquart), *Culex laticinctus* (Edwards) and *Culex pipiens* (Linnaeus). The egg rafts of these three species appeared in similar amounts and consisted of 34.3%, 34%, 31.6%, 25.6%, 40.9%, and 33.5% respectively.

In the first field experiment with signals of crowded *C. longiareolata*, the three species showed significantly different responses to the signals. *Culiseta longiareolata* egg raft distribution was not affected by the treatments (F_2,12_ =0.1; p=0.9). By contrast, *C. laticinctus* oviposition was significantly affected by the different treatments (F_2,12_ =5.25; p=0.02) with significantly fewer egg rafts oviposited in pools containing signals of either high or medium-crowded larvae (Figure 2). The oviposition distribution pattern of *C. pipiens* was not affected by the treatments (F_2,12_ =0.39; p=0.69).

**Figure 2.**
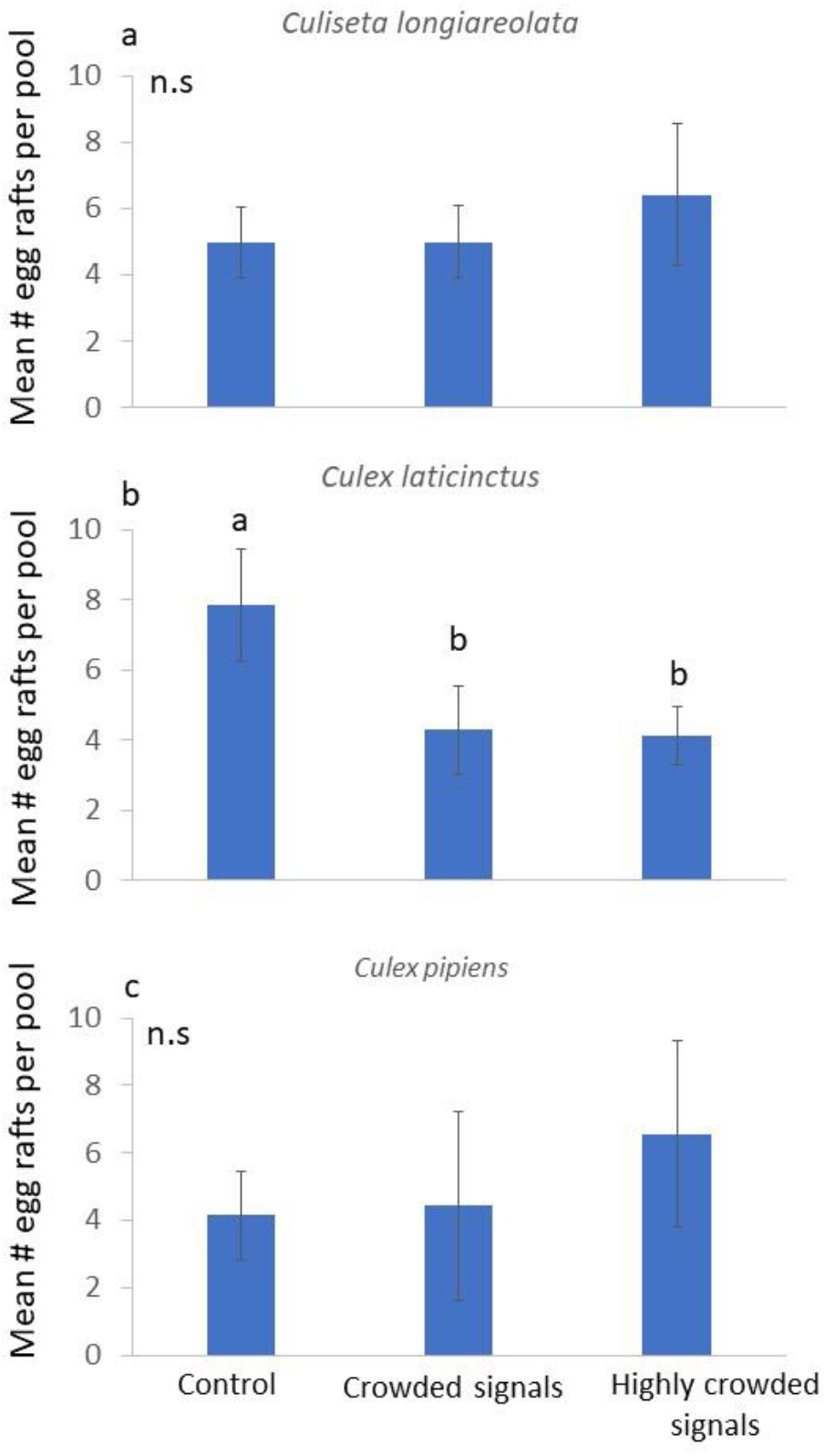
Effect of *Culiseta logiareolata* larvae conditioned water on ovipositing a. *Culiseta logiareolata*; b. *Culex laticinctus*; and c. *Culex pipiens* oviposition, n=7 per treatment, error bars stand for ±1 SE. Different letters indicate treatments that are significantly different based on post hoc comparisons. N.s., not significant.

In the second field experiment with live larvae in the pools, *C. longiareolata* oviposition showed a dramatic response to the presence of larvae (F_2,14_=10.64; p=0.002), with many more egg rafts found in pools added with 200 larvae per pool when compared to pools added with 20 larvae per pool or control pools (Figure 3a). The egg raft distribution of both *C. laticinctus* and *C. pipiens* was not significantly affected by the presence of *C. longiareolata* larvae (F_2,12_=5.25; p=0.87 and F_2,12_ =0.39; p=0.69 respectively, Figure 3b-c).

**Figure 3.**
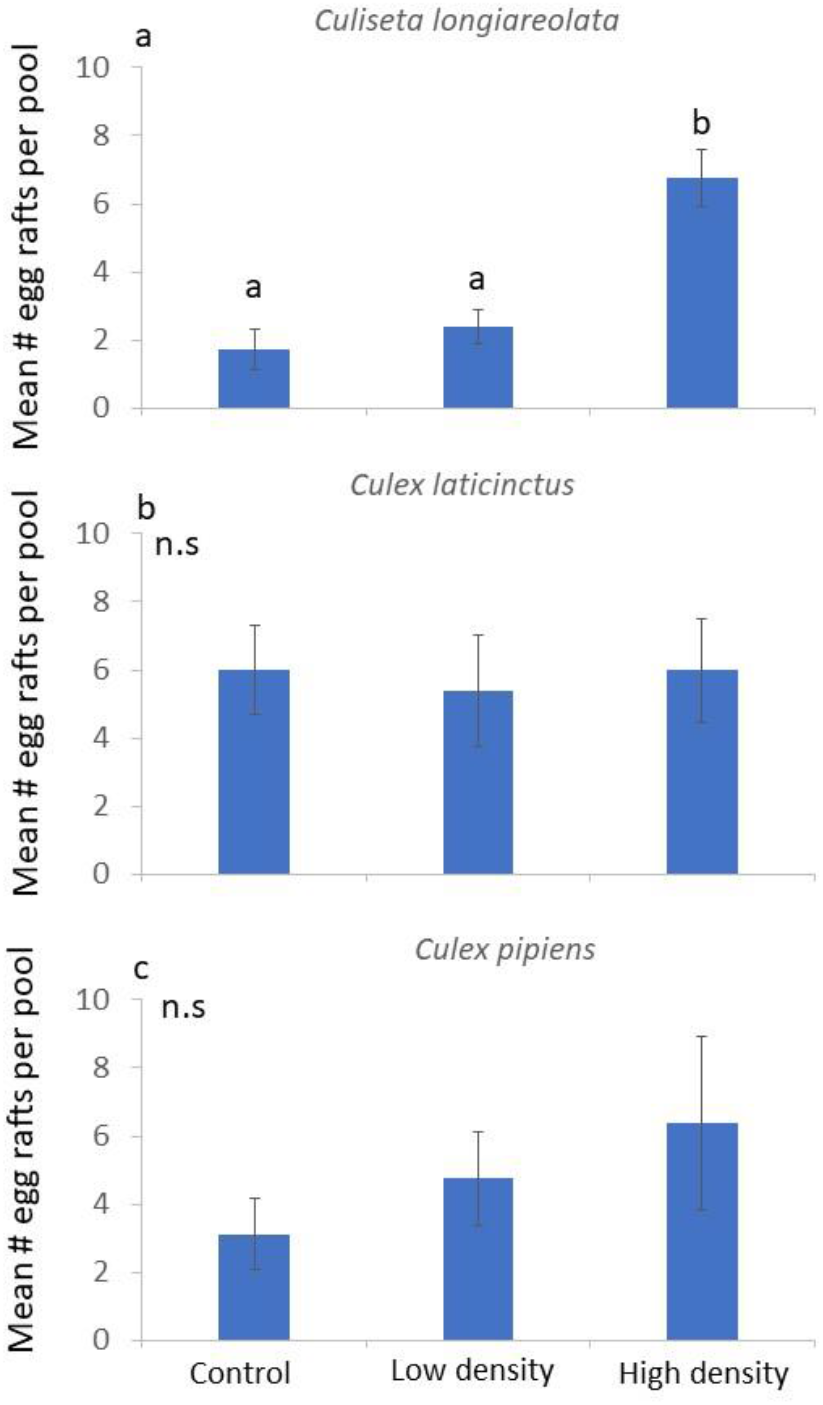
Effect of *Culiseta logiareolata* larvae on ovipositing a. *Culiseta logiareolata*; b. *Culex laticinctus*; and c. *Culex pipiens* oviposition, n=8 per treatment, error bars stand for ±1 SE. Different letters indicate treatments that are significantly different based on post hoc comparisons. N.s., not significant.

## Discussion

This study focused on the role of semiochemicals indicating increasing competition in habitat selection. We hypothesized that water containing highly crowded *C. longiareolata* larvae (20 and 200 larvae in 400 ml.) will contain chemical signals. These signals would be associated with highly crowded larval habitats, and avoided by ovipositing females. *Culiseta longiareolata* females did not respond to these conspecific crowding signals (Figure 2a). Furthermore, a similar number of conspecific larvae that were not crowded were attractive to gravid females (figure 3a). Conspecific density does not necessarily cause an immediate decrease in habitat suitability. A low number of conspecifics may be favorable to colonizers over an empty habitat (Fretwell and Lucas 1969). This trend was shown for ovipositing mosquitoes in response to increasing conspecific larvae (Wasserberg et al. 2014) and eggs (Williams et al. 2008). It is suggested that a habitat with low conspecific density may indicate site persistence, potential mates, and overall appropriate conditions without increased competition.

By contrast to conspecifics, the presence of interspecific-dominant competitors reduces habitat quality even at low densities (Forsman et al. 2008, Eccard and Ylönen 2002, Abramsky et al. 1990). Mosquito larvae are often confined to the oviposition site until metamorphosis. The presence of interspecific larvae from a dominant competing species at this site will often result in reduced survival (Livdahl and Willey 1991). Even if survival to metamorphosis is not significantly reduced, interspecific competition results in other factors associated with a decline in population size as reduced adult body size, longevity, or changes in time to metamorphosis (Silberbush et al. 2014). In our case, competition can be completely avoided by placing the larvae in a competition-free habitat during oviposition. Our results show that females of the subordinate *C. laticinctus* preferred to oviposit in pools that lacked condition water that previously contained crowded larvae of the dominant *C. longiareolata* larvae (Figure 2b). Similar numbers of larvae placed in a higher volume of water did not trigger a significant response (Figure 3b). these results strongly indicate that ovipositing females responded to chemical signals from crowded larvae, indicating a habitat with high competition.

The distribution of *C. pipiens* egg rafts was not significantly affected by the *C. longiareolata* cues (Figure 2c) or larvae (Figure 3c). The global distribution of *C. laticinctus* generally overlaps with that of *C. longiareolata* (Becker et al. 2010). The larvae of these two species often co-occur in recently filled freshwater pools (Kiflawi et al. 2003, Becker et al. 2010, Margalit and Tahori 1974). *Culex pipiens* on the other hand, are characterized by global distribution and by their ability to inhabit a very wide variety of water sources (Becker et al. 2010). As such, this species may be less sensitive to competition and cannot therefore detect and respond to them.

## Acknowledgments

Hatem Abu Raiya and Alon Ornai provided logistical support for the field experiments. Ido Tsurim and Avi Bar-Massada helped with numerous aspects of the study. This work was supported by the Israel Science Foundation Grant number 1166/22 awarded to A. Silberbush and Y. Gerchman. The authors declare no conflict of interest.

